# Paired C-type lectin receptors mediate specific recognition of divergent oomycete pathogens in *C. elegans*

**DOI:** 10.1101/2024.09.05.611528

**Authors:** Kenneth Liu, Manish Grover, Franziska Trusch, Christina Vagena-Pantoula, Domenica Ippolito, Michalis Barkoulas

## Abstract

Innate immune responses can be initiated through the detection of pathogen or damage-associated molecular patterns by host receptors that are often present on the surface of immune cells. While certain invertebrates like *Caenorhabditis elegans* lack professional immune cells, they still respond to infection in a pathogen-specific manner. It has been debated for years whether homologues of the canonical pathogen recognition receptors are also functioning in the nematode. Here we show that C-type lectin receptors mediate species-specific recognition of divergent oomycetes in *C. elegans.* A CLEC-27/CLEC-35 pair is essential for recognition of the oomycete *Myzocytiopsis humicola*, while a CLEC-26/CLEC-36 pair is required for detection of *Haptoglossa zoospora.* Both *clec* pairs are transcriptionally regulated through a shared promoter by the conserved PRD-like homeodomain transcription factor CEH-37/OTX2 and act in sensory neurons and the anterior intestine to trigger a protective immune response in the epidermis. This system enables redundant tissue sensing of oomycete threats through canonical CLEC receptors and host defense via cross-tissue communication.

**Highlights:**

- A CLEC-27/CLEC-35 pair is required for recognition of the oomycete *Myzocytiopsis humicola*
- A CLEC-26/CLEC-36 pair is required for recognition of the oomycete *Haptoglossa zoospora*
- Both CLEC pairs are co-regulated by the homeodomain transcription factor CEH-37/OTX2
- Both CLEC pairs function redundantly in sensory neurons and the intestine for host defense

## Introduction

In their natural habitats, all organisms face a plethora of microbes, therefore it is advantageous to distinguish what is potentially harmful from what is not^1^. To detect pathogens, the innate immune system is equipped with pathogen recognition receptors that sense pathogen-associated molecular patterns and enable animals to either avoid pathogens or combat them with anticipatory immune responses^2^. Such receptors, including the Toll-like receptors (TLR), the nucleotide-binding and oligomerization domain-like receptors, and C-type lectin receptors are found mainly in professional immune cells in mammals^2^. However, many invertebrates such as the nematode *Caenorhabditis elegans* do not have professional immune cells, and instead rely on non-professional cells for defense, such as epithelial cells and neurons, which act together to mount protective immune responses^3–5^. While there is clear evidence that *C. elegans* can sense microbial products^6–9^ and mount defense responses in a pathogen-specific way^10–14^, the mechanisms through which this specificity is achieved remain elusive^15^. This is largely because canonical pattern recognition receptors are either absent or have not been directly linked to immunity in *C. elegans*^16,17^. For example, a single TLR gene named *tol-1* has been shown to modulate the behavioural response to pathogenic bacteria, however, this role is not directly linked to pathogen recognition, differing from how TLRs function in other invertebrates and vertebrates^18,19^. Our understanding of pathogen recognition receptors in *C. elegans* is therefore still in its infancy, with only a few well-established examples reported so far, such as the G protein-coupled receptor DCAR-1 that senses a cuticle damage-induced metabolite produced upon infection with the fungus *Drechmeria coniospora*^20^, the RIG-I homolog DRH-1 that recognises Orsay virus replication products^21^, and the nuclear hormone receptor NHR-86 that directly binds a metabolite from *Pseudomonas aeruginosa* to activate a protective transcriptional program^22^.

One example of a gene family in *C. elegans* that has the potential to encode pattern recognition receptors is the C-type lectin family^23^. In vertebrates, CLEC proteins play fundamental roles in immunity acting either as secreted antimicrobial effectors or as transmembrane pathogen-recognition receptors binding mostly to sugar moieties from a wide range of pathogens^24,25^. However, much less is known about CLEC proteins in invertebrates, where expansion of *clec* gene families can be found^26–29^. In *C. elegans*, the *clec* gene family contains more than 280 members, making it one of the largest gene families in the nematode^30^. While several *clec* genes are differentially expressed upon exposure to different pathogens^23,30–36^, their exact roles in immunity are still poorly understood, and this is particularly pertinent to transmembrane CLEC proteins as most studies have focused on secreted CLEC proteins. Members of the *clec* gene family have been shown to modulate host defense against bacterial pathogens^11,32,36,37,38,39^. In some cases, CLEC proteins display antimicrobial activity and can directly bind to bacterial pathogens^11,36–38,40^. Alternatively, CLEC proteins are also able to modulate immunity in other ways, such as through regulating proteostasis, as shown for CLEC-1 that is secreted by body muscle cells to prevent aggregation of LYS-7 required for fighting against *Bacillus thuringienesis*^41^, or through changes in animal behaviour^33,42^. Given that most C-type lectin domains in *C. elegans* are divergent in sequence in comparison to those found in mammalian proteins^23,30,43^, and that CLEC proteins can play roles beyond immunity (for example in the context of development and ageing^44–46^), it is still unclear whether they act as pattern recognition receptors in the nematode and whether expansion of the *clec* gene family relates to immune response specificity.

Here, we address this question in the context of natural infection of *C. elegans* by two phylogenetically distinct oomycetes^47^, namely *Myzocytiopsis humicola*^12^ and *Haptoglossa zoospora*^48^. Upon oomycete exposure, *C. elegans* activates a specific transcriptional response (ORR – oomycete recognition response) characterised by the epidermal expression of some *chitinase-like (chil)* genes that protect *C. elegans* from oomycete infection^48,49^. We demonstrate that paired CLEC proteins act as plausible receptors for oomycete recognition in *C. elegans,* where a CLEC-27 and CLEC-35 pair specifically recognises *M. humicola*, and a CLEC-26 and CLEC-36 pair specifically recognises *H. zoospora*. Both *clec* pairs are regulated by the conserved PRD-like homeodomain protein CEH-37/OTX2^50^ through a single shared promoter for each pair and predominantly function in the intestine and the AWA sensory neurons, providing a redundant sensing system that leads to epidermal *chil*-27 gene activation through cross-tissue communication. These findings highlight that C-type lectin proteins act as pattern recognition receptors in *C. elegans* and that the diversification of the *clec* gene family contributes to immune specificity.

## Results and Discussion

### *C. elegans* lacking either *ceh-37*, *clec-27* or *clec-35* fail to induce the oomycete recognition response

Through a forward genetic approach aimed at identifying animals that fail to respond to an innocuous *M. humicola* extract that is known to induce the ORR^49^ (Figure S1A), we found independent mutants harbouring premature stop codons within the *clec-27* and *clec-35* genes that resulted in complete loss of *chil-27p::GFP* induction (Figure 1A, B and S1B). CLEC-27 and CLEC-35 belong to the membrane-associated type of C-type lectin proteins^30^. The alleles we recovered carry loss-of-function mutations given that *clec-27* or *clec-35* RNAi led to loss of *chil-27p::GFP* induction upon exposure to *M. humicola* extract (Figure S1C). Furthermore, expression of wild-type *clec-27* and *clec-35* in the respective mutants fully rescued the induction of *chil-27p::GFP* (Figure 1C, S1D). In addition to *clec* genes, we also found non-synonymous changes in the PRD-like homeodomain protein CEH-37^51^ (Figure 1A, B), which has been shown to play roles in sensory neuron specification^50^, telomere function^52^, and intestinal immunity against bacterial pathogens^53^. These mutations also created loss-of-function alleles as *chil-27p::GFP* was not induced upon exposure to *M. humicola* extract when *ceh-37* expression was knocked-down by RNAi (Figure S1C) or in animals carrying a previously described *ceh-37(ok642)* deletion allele^54^ (Figure S1B). Fosmid-based overexpression of the *ceh-37* genetic locus in *ceh-37(icb36)* mutants fully rescued *chil-27* induction upon pathogen extract exposure (Figure 1C, S1D).

**Figure 1:**
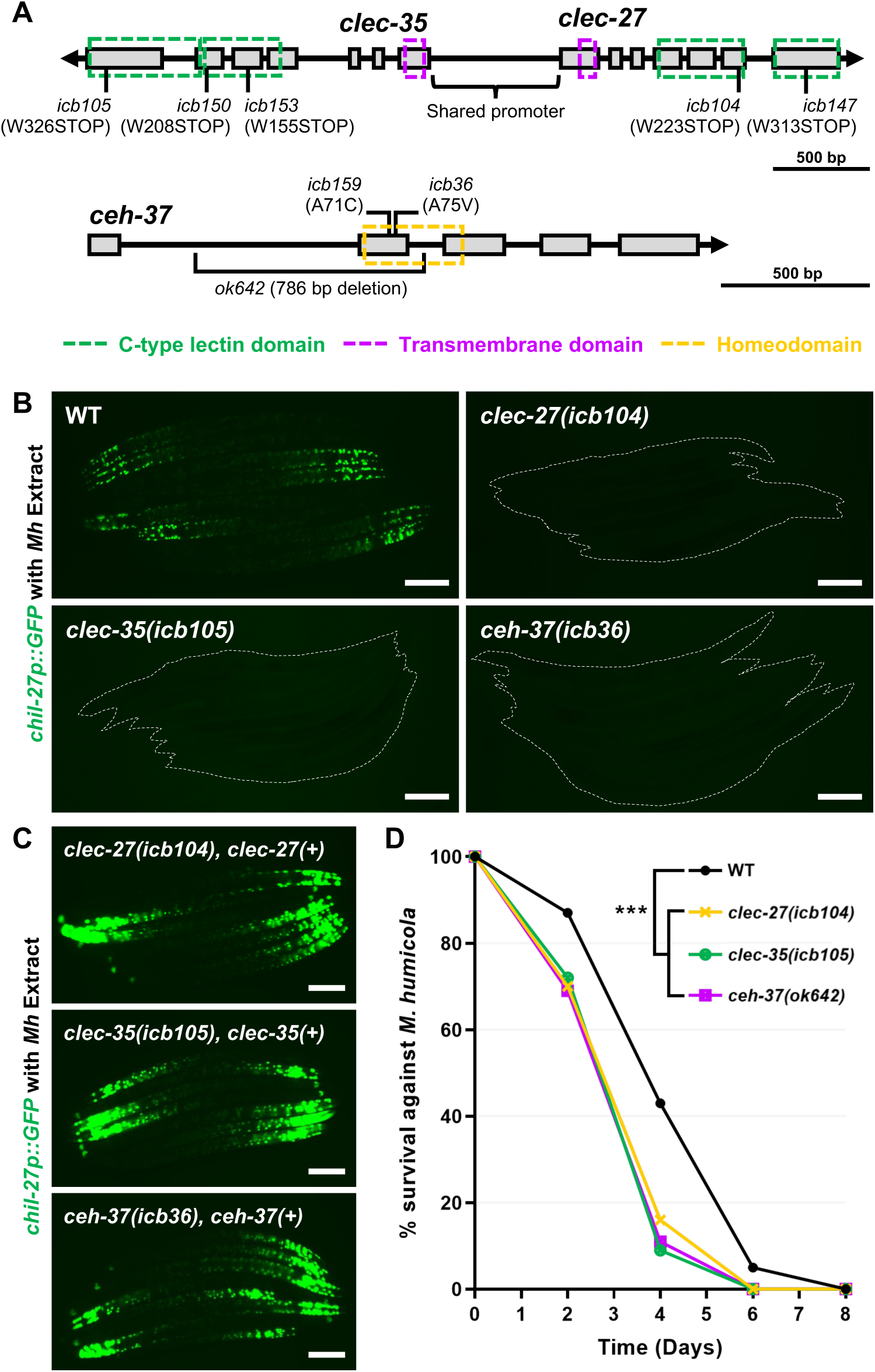
Loss-of-function of *clec-27*, *clec-35* or *ceh-37* impairs the *C. elegans* response to *M. humicola* extract. **(A)** Gene structure of *ceh-37, clec-27 and clec-35* showing the positions of mutant alleles generated in this study. **(B)** L4 stage *clec-27(icb104)*, *clec-35(icb105)* and *ceh-37(icb36)* mutant animals showing lack of *chil-27p::GFP* induction upon exposure to *M. humicola* (*Mh*) extract. **(C)** Rescue of *chil-27p::GFP* induction by providing the wild-type gene locus in the respective mutant backgrounds. **(D)** *clec-27(icb104), clec-35(icb105)* and *ceh-37(ok642)* mutant animals exhibit hypersusceptibility to infection by *M. humicola* (n=60 per condition, performed in triplicates, \*\*\**p<*0.001 based on log-rank test). Scale bars in B, C are 100 µm.

Next, we investigated whether the induction of the oomycete recognition response (ORR) was more broadly impaired in the identified mutants. We performed RNAseq analysis in *clec-27(icb104), clec-35(icb105),* and *ceh-37(ok642)* animals exposed to *M. humicola* extract in comparison to extract-exposed wild-type animals. We found that most of the 50 reproducibly induced genes from the stringent ORR gene list^55^ were not upregulated in the mutants (Figure S1E). Consistent with a defect in mounting the ORR, these mutants were hypersusceptible to infection by *M. humicola* (Figure 1D). Therefore *clec-27*, *clec-35* and *ceh-37* are essential genes required for the induction of the protective ORR in response to *M. humicola* extract exposure.

### *clec-27* and *clec-35* are co-expressed in the neurons and the intestine where they are regulated by CEH-37

Interestingly, *clec-27* and *clec-35* are neighbouring loci on chromosome V and are transcribed in opposite directions, suggesting that they are likely to be co-expressed via a shared promoter (Figure 1A). To test this possibility, we generated a transcriptional reporter where the *clec-27* coding sequence was replaced with GFP and the *clec-35* sequence with mScarlet. Transgenic animals showed GFP and mScarlet colocalization both in neurons in the head area as well as the anterior intestine (Figure 2A). This neuronal and intestinal *clec-27/35* localisation suggested that they may regulate epidermal *chil-27* gene activation in a cross-tissue manner. To address if *clec-27/35* are required in the intestine and/or neurons, we performed tissue-specific rescue experiments in *clec-35(icb105)* mutants. We found that both neuronal (*rab-3p*) and intestinal (*vha-6p*) *clec-35* expression rescued *chil-27p::GFP* induction upon exposure to *M. humicola* extract, whereas epidermal (*dpy-7p*) expression did not (Figure 2B, C). The rescue was also observed at the level of survival in the presence of *M. humicola*, wherein both neuronal and intestinal *clec-27/35* expression reversed the hypersusceptibility of *clec-35(icb105)* mutants (Figure 2D). Both intestine-specific and neuron-enhanced *clec-27* and *clec-35* RNAi significantly reduced *chil-27p::GFP* expression (Figure 2E). Furthermore, CLEC-27 was found to interact with CLEC-35 in a heterologous eukaryotic expression system (Figure S2). These results suggest that *clec-27* and *clec-35* act together in both the neurons and the intestine for oomycete recognition.

**Figure 2:**
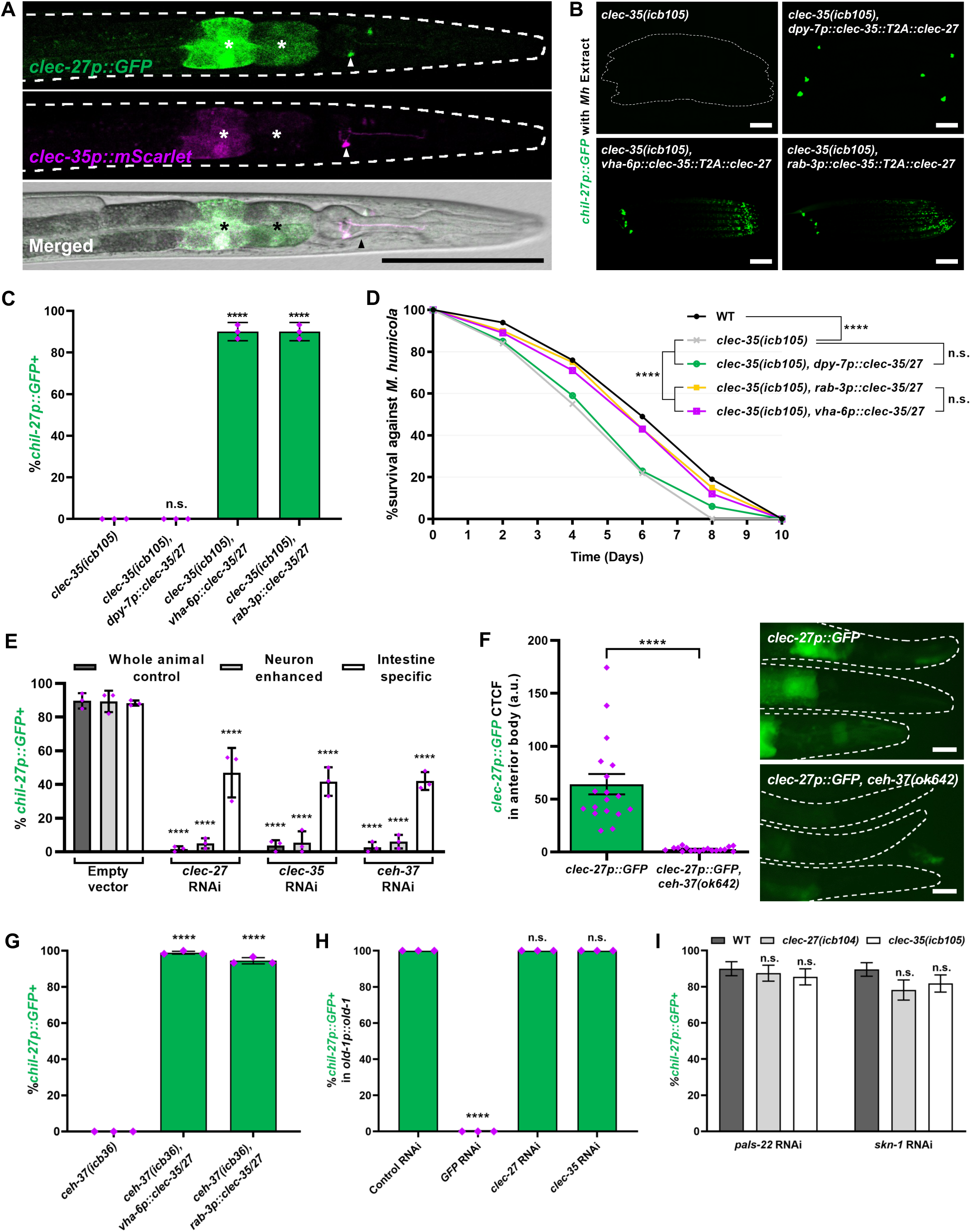
*clec-27* and *clec-35* are co-expressed in neurons and intestine and are regulated by CEH-37. **(A)** Expression of a transcriptional reporter of *clec-27*/*35* (*mScarlet::clec-27/35p::GFP*) shows fluorescence protein co-localisation in the anterior intestine (marked by asterisks) as well as neurons (arrow). **(B and C)** Both neuronal (*rab-3p*) and intestinal (*vha-6p*) expression of *clec-27/35* rescues induction of *chil-27p::GFP* in *clec-35(icb105)* mutants upon exposure to *M. humicola* extract (n=60 animals, performed in triplicates, \*\*\*\**p<*0.0001 based on chi-squared test). **(D)** Neuronal and intestinal expression of *clec-27/35* in *clec-35(icb105)* mutants rescues their hypersusceptibility to *M. humicola* infection, whereas epidermal expression does not (n=60 per condition, performed in triplicate, *\*\*\*\*p<*0.0001 based on log-rank test). **(E)** *chil-27p::GFP* induction upon *M. humicola* extract exposure following tissue-specific RNAi of *clec-27, clec-35,* and *ceh-37* (n=50 per condition, performed in triplicate, *\*\*\*\*p<*0.0001 based on chi-squared test). **(F)** Loss of *clec-27p::GFP* expression in the anterior body in *ceh-37(ok642)* mutants (n=18, ****p<0.0001 based on unpaired t-test). **(G)** Neuronal (*rab-3p*) and intestinal (*vha-6p*) overexpression of *clec-27/35* in *ceh-37(icb36)* mutant rescue *chil-27p::GFP* induction upon exposure to *M. humicola* extract (n=60 animals, performed in triplicates, \*\*\*\**p<*0.0001 based on chi-squared test). **(H)** Quantification of constitutive *chil-27p::GFP* expression upon *old-1* overexpression observed upon *clec-27* and *clec-35* RNAi. RNAi against *gfp* was used as positive control. (n=50 per condition, performed in triplicate, *\*\*\*\*p<*0.0001 based on chi-squared test). **(I)** Quantification of *chil-27p::GFP* induction in *clec-27(icb104)* and *clec-35(icb105)* mutant animals upon *skn-1* and *pals-22* RNAi (n=50 and significance was assessed with chi-squared test). Scale bars in A and B are 100 µm and in F 25 µm. Error bars in C, E, G, H, and I represent standard error of proportion. Error bars in F represent standard error of mean.

Similar to *clec*-*27/35* RNAi, *ceh-37* RNAi in both neuron-enhanced and intestine-specific RNAi strains significantly reduced *chil-27p::GFP* induction (Figure 2E). Notably, we observed in our RNAseq datasets that in uninduced conditions, several *clec* genes, including *clec-27* and *clec-35,* were down-regulated in the *ceh-37(ok642)* mutant background (Table S1). Furthermore, *ceh-37* is expressed in both head neurons^56^ and the intestine^53^ (Figure S3A), and a sequence similar to the consensus sequence required for the DNA binding of the CEH-37 homolog OTX2 in mammals^57^ is present in the shared *clec-27/35* promoter (Figure S3B). Therefore, we reasoned that CEH-37 may regulate the expression of *clec* genes in these tissues. We introduced the *clec-27/35* transcriptional reporter in *ceh-37(ok642)* deletion mutant and found significant reduction of *clec-27p::GFP* expression in the anterior intestine and in the head region (Figure 2F). Furthermore, restoring *clec-27*/*35* expression in neurons (*rab-3p*) or intestine (*vha-6p*) in *ceh-37(icb36)* mutants was sufficient to rescue *chil-27p::GFP* induction upon exposure to *M. humicola* extract (Figure 2G). Taken together, these results suggest that the loss of *chil-27p::GFP* induction upon oomycete recognition in *ceh-37* mutants is because CEH-37 regulates the expression of *clec-27/35* in both neurons and the intestine.

We have previously shown that ORR activation can also be triggered through perturbations directly in the epidermis, such as proteasome dysfunction^58^, modulation of the PALS-22/25 module^59,60^, or constitutive signalling by the receptor tyrosine kinase OLD-1^55^. We hypothesised that CLEC-27/35 are likely to act prior to the activation of immune response in the epidermis. To test this hypothesis, we performed *clec-27/35* RNAi in animals showing constitutive expression of *chil-27p::GFP* due to overexpression of *old-1* (Figure 2H) or, conversely, *pals-22* and *skn-*1 RNAi in the *clec-27/35* mutant backgrounds recovered from our genetic screen (Figure 2I). In all cases, we observed normal *chil-27p::GFP* induction, confirming that the role of *clec-27/35* in the neurons and intestine lies upstream of the epidermal triggers of the ORR.

### AWA is the main oomycete sensing neuron in *C. elegans*

To identify the specific neurons involved in *M. humicola* extract recognition, we focused on AWA, AWB, ASG and BAG, which are the four neurons that express CEH-37^56^. We attempted to rescue c*lec-27(icb104)* mutants by restoring *clec-27* under the neuron-specific promoters *odr-7p* (AWA)^61^, *str-1p* (AWB)^62^, *gcy-15p* (ASG)^63^ and *gcy-33p* (BAG)^64^. As controls, we restored *clec-27* expression in *clec-27(icb104)* mutants under the *ceh-37* promoter in case multiple neurons are required, or under the *srg-8p* promoter (ASK)^65^, where *ceh-37* is not normally expressed^56^. Here we found that *clec-27/35* rescue in the AWA neurons most strongly recovered *chil-27p::GFP* induction upon exposure to *M. humicola* extract followed by rescue in AWB neurons (Figure 3A). Furthermore, restoring *clec-27/35* expression in *ceh-37(icb36)* mutants in either AWA or AWB neurons was sufficient for animals to strongly recover *chil-27p::GFP* induction upon exposure to *M. humicola* extract (Figure 3B).

**Figure 3:**
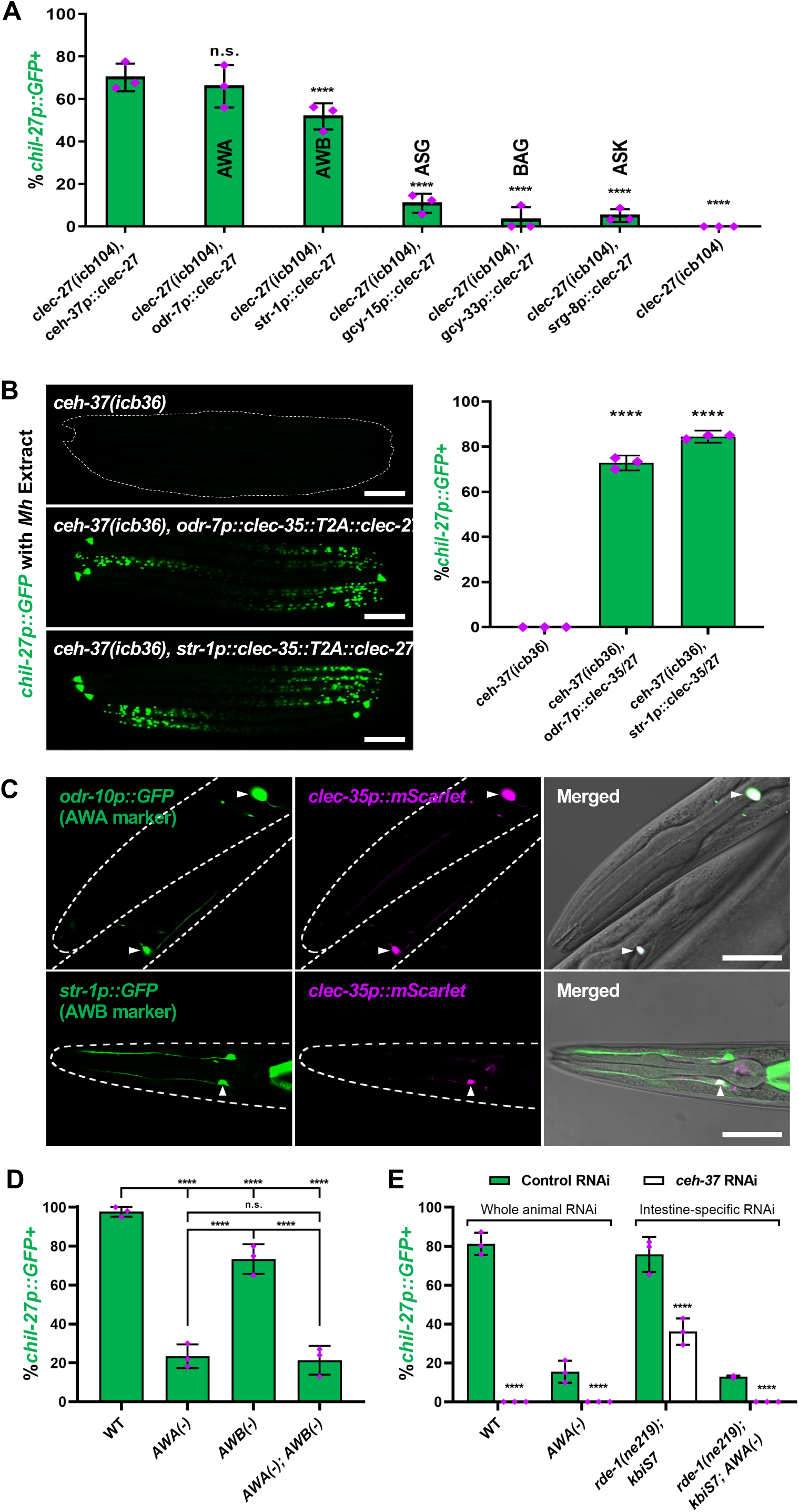
AWA is the main oomycete sensing neuron. **(A)** Rescue of *chil-27p::GFP* induction in *clec-27(icb104)* mutants upon *M. humicola* extract treatment by restoring *clec-27* expression in specific neurons. Note that *clec-27* expression in AWA neurons most strongly rescues induction of *chil-27p::GFP* upon *M. humicola* extract exposure, followed by *clec-27* expression in AWB neurons (n>60 animals, performed in triplicates, \*\*\*\**p<*0.0001 based on Cochran–Mantel–Haenszel test). **(B)** AWA (*odr-7p*) and AWB (*str-1p*) specific expression of *clec-27/35* in *ceh-37(icb36)* is sufficient to rescue induction of *chil-27p::GFP* upon *M. humicola* extract treatment (n=60 animals, performed in triplicates, \*\*\*\**p<*0.0001 based on chi-squared test). **(C)** Co-localisation of *clec-35p::mScarlet* with AWA (*odr-10p::GFP*) and AWB (*str-1p::GFP*) specific markers confirms expression of *clec-27/35* in these neurons (shown with arrows). **(D)** Genetic ablation of AWA neurons strongly reduces *chil-27p::GFP* induction upon exposure to *M. humicola* extract compared to ablation of AWB neurons (n=50 animals, performed in triplicates, \*\*\*\**p<*0.0001 based on chi-squared test). **(E)** AWA ablation combined with intestine-specific RNAi of *ceh-37* leads to complete loss of *chil-27p::GFP* expression (n=50 animals, performed in triplicates, \*\*\*\**p<*0.0001 based on chi-squared test). Scale bar is 100 µm in B and 25 µm in C. Error bars in A represent standard deviation. Error bars in B, D, and E represent standard error of proportion.

To confirm that *clec-27/35* are expressed in these neurons we co-localised the neuron-specific markers *odr-10p::GFP* (AWA)^66^ and *str-1p::GFP* (AWB) with *clec-27/35p::mScarlet*. We found *clec-35* expression in both AWA and AWB neurons (Figure 3C). To test the functional significance of the AWA/AWB neurons, we genetically ablated them by cell-specific expression of a caspase and assessed whether animals can mount *chil-27p::GFP* expression upon *M. humicola* extract exposure. While ablation of AWA neurons^67^ showed strong reduction of *chil-27p::GFP* upon exposure to *M. humicola* extract, AWB ablation^68^ caused a milder effect. Simultaneous ablation of both neurons did not result in a further reduction in response compared to AWA ablation alone (Figure 3D). We note that the partial *chil-27p::GFP* induction observed in the AWA-ablated strain is likely due to the intestinal expression of *clec-27/35* driving recognition, as simultaneous AWA ablation and intestinal-specific *ceh-37* RNAi resulted in complete loss of *chil-27p::GFP* induction (Figure 3E). This suggests that while AWB neurons possess the capability of contributing to oomycete recognition, neuronal oomycete recognition is predominantly mediated by the AWA neurons.

### A different CLEC receptor pair in AWA neurons mediates recognition of the phylogenetically divergent oomycete *Haptoglossa zoospora*

It is known that *C. elegans* is able to sense oomycetes that are phylogenetically distinct from *M. humicola,* such as those belonging to the early diverging *Haptoglossa* genus^48^. Therefore, we examined if known regulators of the response to *M. humicola* extract^55^ were also relevant for *Haptoglossa zoospora* extract recognition. Surprisingly, we observed strong induction of *chil-27p::GFP* in both *clec-27* and *clec-35* mutants, while there was a strong loss of *chil-27p::GFP* induction in *ceh-37(ok642)*, *old-1(ok1273), flor-1(icb116), and vab-3(icb127)* mutants (Figure 4A). Hence, CLEC-27 and CLEC-35 are not required for *H. zoospora* recognition, however, the required receptor(s) are likely regulated by CEH-37.

**Figure 4:**
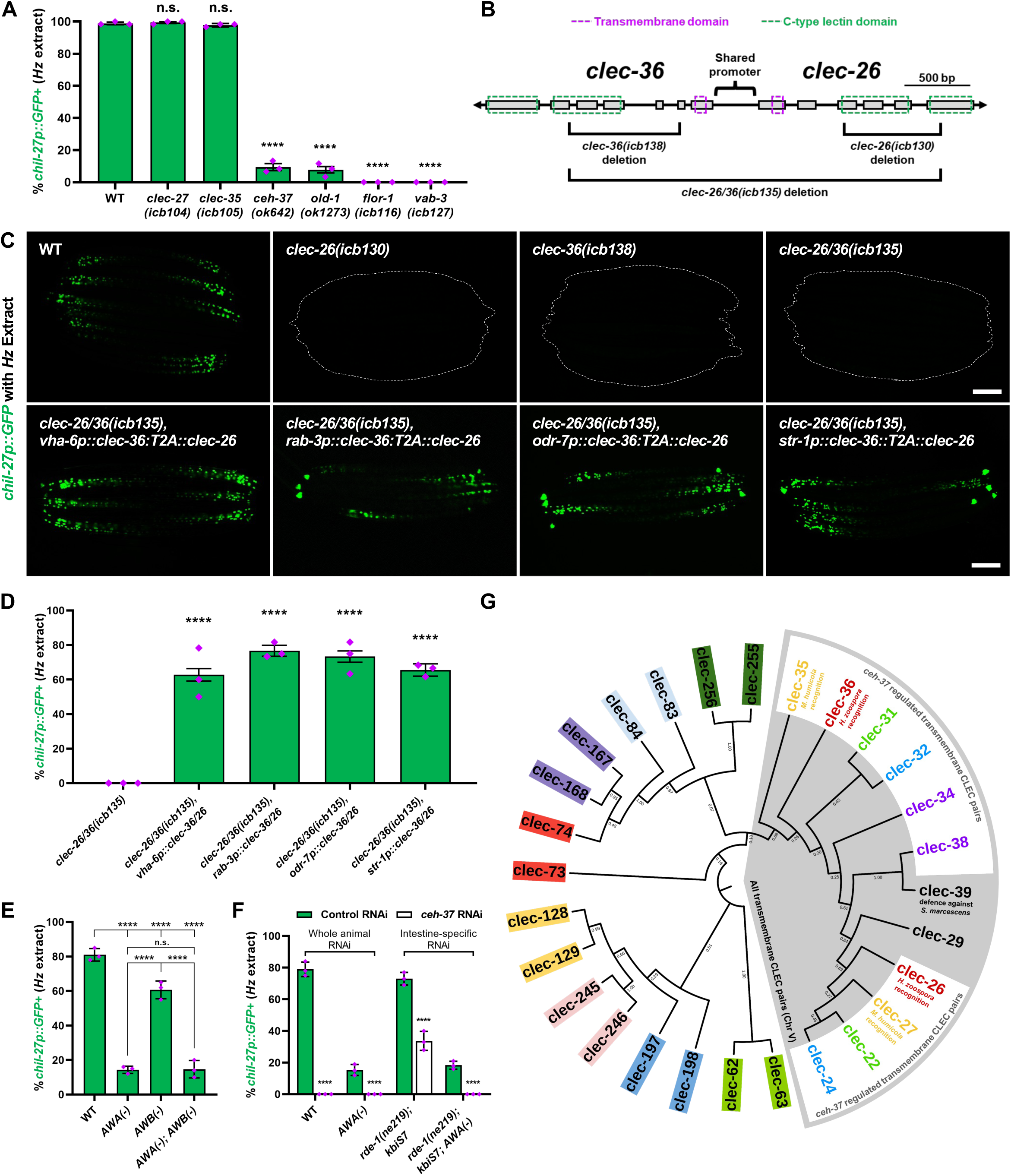
A CLEC-26/36 pair mediates recognition of the oomycete Haptoglossa zoospora. **(A)** Induction of *chil-27p::GFP* in *clec-27(icb104), clec-35(icb105)*, *ceh-37(ok642), old-1(ok1273), flor-1(icb116),* and *vab-3(icb127)* mutants upon exposure to *H. zoospora* (Hz) extract as compared to wild-type animals (n=60 per condition, performed in triplicate, *\*\*\*\*p<*0.0001 based on chi-squared test). **(B)** Genomic organization of *clec-26* and *clec-36* loci and mutations generated in the study. **(C)** Neuronal (*rab-3p*), intestinal (*vha-6p*), AWA-specific (*odr-7p*), and AWB-specific (*str-1p*) expression of *clec-26/36* are sufficient to rescue *chil-27p::GFP* induction in *clec-26/36(icb135)* mutants upon exposure to *H. zoospora* extract. **(D)** Quantification of *chil-27p::GFP* induction in the rescued lines above (n=60 per condition, performed in triplicate, *\*\*\*\*p<*0.0001 based on chi-squared test). **(E)** Genetic ablation of AWA and less so of AWB neurons compromises induction of *chil-27p::GFP* upon exposure to *H. zoospora* extract (n=50 per condition, performed in triplicate, *\*\*\*\*p<*0.0001 based on chi-squared test). **(F)** Genetic ablation of AWA combined with intestine-specific knockdown of *ceh-37* significantly compromises induction of *chil-27p::GFP* upon exposure to *H. zoospora* extract (n=60 per condition, performed in triplicate, *\*\*\*\*p<*0.0001 based on chi-squared test). **(G)** Maximum-likelihood phylogenetic tree with 500 bootstrap replications showing the relationship between the *clec-27/35* and *clec-26/36* pairs. All paired protein-coding *clec* genes (indicated by the same colour) with predicted bidirectional promoters present in the *C. elegans* genome are shown. All *clec* pairs encoding transmembrane proteins are shown within the grey sector, those that are regulated by CEH-37 are further highlighted in white within the grey sector. Scale bar in C is 100 µm. Error bars in A, D, E, and F represent standard error of proportion.

To identify if CEH-37-regulated *clecs* are required for *H. zoospora* extract recognition, we performed a targeted RNAi screen focusing on *clec* genes that were found to be downregulated in *ceh-37(ok642)* mutants via RNAseq (Table S1). In this screen, L4 stage animals carrying the *chil-27p::GFP* reporter were subjected to RNAi targeting CEH-37-regulated *clec* genes before *H. zoospora* extract was added to the plates. Interestingly, only RNAi of *clec-26* and *clec-36* resulted in near complete loss of *chil-27p::GFP* induction upon 48 hours of exposure to *H. zoospora* extract (Figure S4A). To consolidate this finding, we generated *clec-26(icb130)*, *clec-36(icb138)*, and *clec-26/36(icb135)* deletion mutants using CRISPR and found loss of *chil-27p::GFP* induction upon exposure to *H. zoospora* extract (Figure 4B, C). Intestinal and neuronal expression of *clec-26/36* in *clec-26/36(icb135)* mutants rescued *chil-27p::GFP* induction upon exposure to *H. zoospora* extract (Figure 4C, D). Consistent with this result, both neuron-enhanced and intestine-specific RNAi of *clec-26*, *clec-36* and *ceh-37* strongly reduced *chil-27p::GFP* induction upon exposure to *H. zoospora* extract (Figure S4B).

Like *clec-27/35*, *clec-26* and *clec-36* are neighbouring loci on chromosome V and are transcribed in opposite directions, thus it is likely that they are co-expressed and co-regulated (Figure 4B). A *clec-26/36* transcriptional reporter generated using the shared *clec-26/36* promoter, which also harbours a sequence resembling the OTX2 binding site (Figure S3B), showed GFP and mScarlet co-localisation in the head region as well as the intestine (Figure S4C), and CLEC-26 was found to interact with CLEC-36 in a heterologous eukaryotic expression system (Figure S2). We also observed co-localization of the *clec-26/36p::mScarlet* transcriptional reporter with the AWA (*odr-10p::GFP*) and AWB (*str-1p::GFP*) neuronal markers (Figure S4D). Rescue of *clec-26/36* expression in AWA or AWB neurons was sufficient to restore *chil-27p::GFP* induction in *clec-26/36(icb135)* mutants upon exposure to *H. zoospora* extract (Figure 4C, D). Compared to wild-type animals, AWA-ablated animals displayed a strong reduction in *chil-27p::GFP* induction upon exposure to *H. zoospora* extract, whereas AWB-ablated animals showed a smaller reduction in *chil-27p::GFP* induction (Figure 4E). Similar to *M. humicola* extract recognition, ablation of both AWA and AWB did not result in further reduction of *chil-27p::GFP* induction compared to AWA ablation alone, suggesting that it is the AWA neurons that primarily mediate neuronal recognition of *H. zoospora*-derived extract (Figure 4E). When genetic ablation of AWA was combined with intestine-specific knockdown of *ceh-37*, we observed complete loss of *chil-27p::GFP* induction upon exposure to *H. zoospora* extract (Figure 4F), highlighting that CLEC-26/36 are likely to act together in sensory neurons and the intestine similar to the CLEC-27/35 pair.

To conclude, our findings demonstrate that selected members of the expanded C-type lectin family in *C. elegans*, physically located next to each other on the genome, can act in pairs to mediate species-specific recognition of divergent oomycete pathogens. Considering the paired genomic organization and the essential requirement for both components of the pair for oomycete recognition, it is plausible that these CLEC proteins function as heterodimeric receptors. It is believed that heterodimerization of CLEC receptors in mammals can increase both the sensitivity for ligands and range of molecules that can be sensed^69^. Our work establishes a paradigm for oomycete recognition wherein canonical pattern recognition receptors act redundantly in non-specialised epithelial and neuronal cells to mount a protective immune response in the skin through cross-tissue communication. Such non-cell autonomous regulation of immunity highlights the complex ways that *C. elegans* can use to sense and respond to its natural eukaryotic pathogens and extends previous reports demonstrating neuronal regulation of host immunity against bacterial infections in *C. elegans*^7,39,70–72^. Activation of CLEC pairs may follow the binding of distinct oomycete-derived ligands and lies upstream of OLD-1/FLOR-1-mediated signalling^55^ which leads to induction of *chil* genes in the epidermis. Interestingly, both the CLEC pair and the OLD-1/FLOR-1 module are regulated by homeodomain proteins (CEH-37 and VAB-3 respectively) known to play broader developmental roles in patterning and sensory organ development^50,73^, and all genes involved in sensing are not induced during infection^12^. We note that among the 283 annotated *clec* genes in the *C. elegans* genome, 28 *clec* genes exist in similar pairs sharing a putative bidirectional promoter, 10 of which are also regulated by CEH-37 (Figure 4G). We therefore speculate that other CLEC proteins acting alone or in pairs are likely to play a role in immune response activation against other pathogens, similar to their canonical pattern recognition role in immune detection in mammals^24^.

## STAR METHODS

Detailed methods are provided in the online version of this paper and include the following:

- KEY RESOURCES TABLE
- RESOURCE AVAILABILITY

- Lead Contact
- Materials Availability
- Data and Code Availability
- EXPERIMENTAL MODEL AND SUBJECT DETAILS
- METHOD DETAILS

- EMS mutagenesis
- Oomycete Infection Assays
- RNAseq
- Microscopy
- Molecular cloning and transgenesis
- CRISPR-mediated genome editing
- RNA Interference (RNAi)
- Single molecule fluorescence in situ hybridization (smFISH)
- Phylogenetic analysis
- Heterologous *clec* expression and co-immunoprecipitation
- QUANTIFICATION AND STATISTICAL ANALYSIS

### Key Resources Table

| REAGENT or RESOURCE | SOURCE | IDENTIFIER |
| --- | --- | --- |
| **Antibodies** |  |  |
| α-GFP, rabbit | Invitrogen | Cat# A11122 |
| α-myc, rat | Proteintech | Cat# 9E1 |
| HRP-coupled antibody, α-rabbit | Merck | Cat# A9169 |
| HRP-coupled antibody, α-rat | Merck | Cat# A9037 |
| **Bacterial and virus strains** |  |  |
| *E. coli* OP50 | Caenorhabditis Genetics Center | Wormbase ID: OP50 |
| Ahringer RNAi Library in *E. coli* HT115 | Source Bioscience | N/A |
| *C. elegans* ORFeome (Vidal RNAi Library) in *E. coli* HT115 | Horizon Discovery | N/A |
| *A. tumefaciens* GV3101 | Tolga Bozkurt Lab Collection (Yuen et al., 2024)<sup>74</sup> | N/A |
| **Chemicals, peptides, and recombinant proteins** |  |  |
| Ethyl methanesulfonate (EMS) | Sigma | Ca# M0880 |
| Isopropyl β-D-1-thiogalactopyranoside (IPTG) | Thermo Scientific | Cat# R0392 |
| TRIzol | Invitrogen | Cat# 15596026 |
| **Critical commercial assays** |  |  |
| QIAquick PCR Purification Kit | Qiagen | Cat# 28106 |
| PureLink Quick Gel Extraction Kit | Invitrogen | Cat# K210012 |
| PureLink Quick Plasmid Miniprep Kit | Invitrogen | Cat# K210011 |
| **Deposited data** |  |  |
| RNA-seq data, normalized per gene sequence counts and differentially expressed genes as determined by Kallisto & Sleuth respectively | This Paper | GEO: GSE263858 |
| **Experimental models: Organisms/strains** |  |  |
| *Myzocytiopsis humicola* (Lisbon Isolate) | Osman et al., 2018<sup>12</sup> | JUo1 |
| *Haptoglossa zoospora* (Eastbourne Isolate) | Grover et al., 2021<sup>48</sup> | MBAo1 |
| *Nicotiana benthamiana* (wildtype) | Tolga Bozkurt Lab<br>Collection (Yuen et al.,<br>2024)<sup>74</sup> | N/A |
| *C. elegans*: wild-type N2 | Caenorhabditis<br>Genetics Center | Wormbase ID: N2 |
| *C. elegans*: *icbIs4*[pGO4, pCMH1195] II | Osman et al., 2018<sup>12</sup> | MBA281 |
| *C. elegans*: *icbIs5*[pGO4, pCMH1195] IV | Osman et al., 2018<sup>12</sup> | MBA282 |
| *C. elegans*: *ceh-37(ok642)* X | Caenorhabditis<br>Genetics Center | LJ2 |
| *C. elegans*: *icbIs4*[pGO4, pCMH1195] II; *ceh-37(icb36)* X | This study | MBA503 |
| *C. elegans*: *icbIs4*[pGO4, pCMH1195] II; *ceh-37(ok642)* X | This study | MBA1054 |
| *C. elegans*: *icbIs4*[pGO4, pCMH1195] II; *ceh-37(icb159)* X | This study | MBA1700 |
| *C. elegans*: *icbIs4*[pGO4, pCMH1195] II; *ceh-37(icb152)* X | This study | MBA1702 |
| *C. elegans*: *icbIs4*[pGO4, pCMH1195] II; *clec-35(icb153)* V | This study | MBA1703 |
| *C. elegans*: *icbIs4*[pGO4, pCMH1195] II; *clec-35(icb150)* V | This study | MBA1697 |
| *C. elegans*: *icbIs4*[pGO4, pCMH1195] II; *clec-35(icb105)* V | This study | MBA1105 |
| *C. elegans*: *icbIs4*[pGO4, pCMH1195] II; *clec-27(icb104)* V | This study | MBA1104 |
| *C. elegans*: *icbIs4*[pGO4, pCMH1195] II; *clec-27(icb147)* V | This study | MBA1693 |
| *C. elegans*: *icbIs4*[pGO4, pCMH1195] II; *clec-27(icb104)* V; *icbEx293[clec-27p::clec-27, bus-1p::GFP, pBJ36]* | This study | MBA1116 |
| *C. elegans*: *icbIs4*[pGO4, pCMH1195] II; *clec-35(icb105)* V; *icbEx296[clec-35p::clec-35, bus-1p::GFP, pBJ36]* | This study | MBA1119 |
| *C. elegans*: *icbEx636[ceh-37(WRM065bE05), bus-1p::GFP, pBJ36]*, *icbIs4*[pGO4, pCMH1195] II; *ceh-37(icb36)* X | This study | MBA1785 |
| *C. elegans*: *clec-27(icb104)* V | This study | MBA1130 |
| *C. elegans*: *clec-35(icb105)* V | This study | MBA1111 |
| *C. elegans*: *icbEx306[pCV12(clec-35 3'UTR::wrmscarlet::clec-35/27p::GFPo::clec-27 3'UTR), bus-1p::GFP]* | This study | MBA1199 |
| *C. elegans*: *icbIs4*[pGO4, pCMH1195] II; *clec-35(icb105)* V; *icbEx634[pCV10(rab-3p::clec-35::T2A::clec-27), bus-1p::GFP, pBJ36]* | This study | MBA1208 |
| *C. elegans*: *icbIs4*[pGO4, pCMH1195] II; *clec-35(icb105)* V; *icbEx635[pCV11(dpy-7p::clec-35::T2A::clec-27), bus-1p::GFP, pBJ36]* | This study | MBA1783 |
| *C. elegans*: *icbIs4*[pGO4, pCMH1195] II; *clec-35(icb105)* V; *icbEx658[pKL158(vha-6p::clec-35::T2A::clec-27), bus-1p::GFP, pBJ36]* | This study | MBA1877 |
| *C. elegans*: *ceh-37(ok642)* X; *icbEx306[pCV12(clec-35 3'UTR::wrmscarlet::clec-35/27p::GFPo::clec-27 3'UTR), bus-1p::GFP, pBJ36]* | This study | MBA1906 |
| *C. elegans*: *icbIs4*[pGO4, pCMH1195] II; *ceh-37(icb36)* X; *icbEX686[pKL158(vha-6p::clec-35::T2A::clec-27), bus-1p::GFP, pBJ36]* | This study | MBA1918 |
| *C. elegans*: *icbEx634[pCV10(rab-3p::clec-35::T2A::clec-27), bus-1p::GFP, pBJ36]*, *icbIs4*[pGO4, pCMH1195] II; *ceh-37(icb36)* X | This study | MBA1784 |
| *C. elegans*: *icbIs4*[pGO4, pCMH1195] II; *clec-27*(*icb104*) V; *icbEx329*[pKL1(*srg-8p::clec-27*), *bus-1p::GFP*, pBJ36] | This study | MBA1298 |
| *C. elegans*: *icbIs4*[pGO4, pCMH1195] II; *clec-27*(*icb104*) V; *icbEx330*[pKL2(*gcy-15p::clec-27*), *bus-1p::GFP*, pBJ36] | This study | MBA1232 |
| *C. elegans*: *icbIs4*[pGO4, pCMH1195] II; *clec-27*(*icb104*) V; *icbEx331*[pKL3(*gcy-33p::clec-27*), *bus-1p::GFP*, pBJ36] | This study | MBA1233 |
| *C. elegans*: *icbIs4*[pGO4, pCMH1195] II; *clec-27*(*icb104*) V; *icbEx332*[pKL4(*str-1p::clec-27*), *bus-1p::GFP*, pBJ36] | This study | MBA1234 |
| *C. elegans*: *icbIs4*[pGO4, pCMH1195] II; *clec-27*(*icb104*) V; *icbEx337*[pKL6(*ceh-37p::clec-27*), *bus-1p::GFP*, pBJ36] | This study | MBA1301 |
| *C. elegans*: *icbIs4*[pGO4, pCMH1195] II; *clec-27*(*icb104*) V; *icbEx339*[pKL7(*odr-7p::clec-27*), *bus-1p::GFP*, pBJ36] | This study | MBA1302 |
| *C. elegans*: *icbIs4*[pGO4, pCMH1195] II; *ceh-37*(*icb36*) X; *icbEx345*[pKL8(*odr-7p::clec-35::T2A::clec-27*), *bus-1p::GFP*, pBJ36] | This study | MBA1306 |
| *C. elegans*: *icbIs4*[pGO4, pCMH1195] II; *ceh-37*(*icb36*) X; *icbEx595*[pKL128(*str-1p::clec-35::T2A::clec-27*), *bus-1p::GFP*, pBJ36] | This study | MBA1732 |
| *C. elegans*: *icbEx450*[pKL53(*odr-10p::GFP*), *PIP1303F*(*rps-0::hgb:unc-54*), pBJ36]; *icbEx515*[pKL71(*clec-35p::wrmscarlet*), *bus-1p::GFP*, pBJ36] | This study | MBA1864 |
| *C. elegans*: *icbEx451*[pKL54(*str-1p::GFP*), *PIP1303F*(*rps-0::hgb:unc-54*), pBJ36]; *icbEx515*[pKL71(*clec-35p::wrmScarlet*), *bus-1p::GFP*, pBJ36] | This study | MBA1598 |
| *C. elegans*: *icbIs4*[pGO4, pCMH1195] II; *Ex[odr-10::mCasp1, myo-3::GFP]* | This study | MBA1754 |
| *C. elegans*: *icbIs4*[pGO4, pCMH1195] II; *pels1715*[*str-1::mCasp1 + unc-122::GFP*]; *Ex[odr-10::mCasp1, myo-3::GFP]* | This study | MBA1755 |
| *C. elegans*: *pels1715*[*str-1p::mCasp-1, unc-122p::GFP*], *icbIs4*[pGO4, pCMH1195] II | Fasseas et al., 2021<sup>49</sup> | MBA1196 |
| *C. elegans*: *kbls7*[*nhx-2p::rde-1*]; *rol-6*(*su1006*) II; *icbIs5*[pGO4, pCMH1195] IV; *rde-1*(*ne219*) V | Grover et al., 2024<sup>58</sup> | MBA597 |
| *C. elegans*: *kbls7*[*nhx-2p::rde-1*]; *rol-6*(*su1006*) II; *icbIs5*[pGO4, pCMH1195] IV; *rde-1*(*ne219*) V; *Ex[odr-10::mCasp1 + myo-3::GFP]* | This study | MBA1907 |
| *C. elegans*: *old-1*(*ok1273*) II; *icbIs4*[pGO4, pCMH1195] IV | Drury et al., 2023<sup>55</sup> | MBA1018 |
| *C. elegans*: *icbIs4*[pGO4, pCMH1195] IV; *flor-1*(*icb116*) V | Drury et al., 2023<sup>55</sup> | MBA1415 |
| *C. elegans*: *icbIs22*[*old-1p::old-1, bus-1p::GFP*, pBJ36] IV; *icbIs5*[pGO4, pCMH1195] IV; *vab-3*(*icb127*) X | Drury et al., 2023<sup>55</sup> | MBA1339 |
| *C. elegans*: *icbIs4*[pGO4, pCMH1195] II; *clec-26*(*icb130*) V | This study | MBA1472 |
| *C. elegans*: *icbIs4*[pGO4, pCMH1195] II; *clec-36*(*icb138*) V | This study | MBA1605 |
| *C. elegans*: *icbIs4*[pGO4, pCMH1195] II; *clec-26/36*(*icb135*) V | This study | MBA1578 |
| *C. elegans*: *icbIs4*[pGO4, pCMH1195] II; *clec-26/36*(*icb135*) V; *icbEx619*[pKL136(*rab-3p::clec-36::GSG-T2A::clec-26*), *bus-1p::GFP*, pBJ36] | This study | MBA1791 |
| *C. elegans*: *icbIs4*[pGO4, pCMH1195] II; *clec-26/36* (*icb135*) V; *icbEx505*[pKL63(*odr-7p::clec-36::GSG-T2A::clec-26*), *bus-1p::GFP*, pBJ36] | This study | MBA1577 |
| *C. elegans*: *icbIs4*[pGO4, pCMH1195] II; *clec-26/36*(*icb135*) V; *icbEx597*[pKL130(*str-1p::clec-36::T2A::clec-26*), *bus-1p::GFP*, pBJ36] | This study | MBA1734 |
| *C. elegans*: *icbIs4*[pGO4, pCMH1195] II; *clec-26/36* (*icb135*) V; *icbEx624*[pKL141(*vha-6p::clec-36::T2A::clec-26*), *bus-1p::GFP*, pBJ36] | This study | MBA1796 |
| *C. elegans*: *icbIs4*[pGO4, pCMH1195] II; *sid-1*(*pk3321*) *uls69* V | Grover et al., 2024<sup>58</sup> | MBA501 |
| *C. elegans*: *icbIs5*[pGO4, pCMH1195] IV; *rde-1*(*ne219*) V; *xkIs99* [*wrt-2p::rde-1::unc-54 3'UTR*] | Drury et al., 2023<sup>55</sup> | MBA576 |
| *C. elegans*: *ceh-37*(*ot1023*[*ceh-37::GFP::FLAG*]) X. | Caenorhabditis Genetics Center | OH16345 |
| *C. elegans*: *icbIs22*[*old-1p::old-1::GFP*, *bus-1p::GFP*, pBJ36] IV; *icbIs5*[pGO4, pCMH1195] IV; | Drury et al., 2023<sup>55</sup> | MBA1437 |
| *C. elegans*: *icbEx449* [pKL52(*wrmScarlet::clec-26/36p::GFP*), *bus-1p::GFP*, pBJ36] | This study | MBA1477 |
| *C. elegans*: *icbEx450* [pKL53(*odr-10p::GFP*), *PIP1303F*(*rps-0::hgb:unc-54*), pBJ36]; *icbEx516*[pKL72(*clec-36p::wrmScarlet*), *bus-1p::GFP*, pBJ36]; | This study | MBA1599 |
| *C. elegans*: *icbEx450* [pKL54(*str-1p::GFP*), *PIP1303F*(*rps-0::hgb:unc-54*), pBJ36]; *icbEx516*[pKL72(*clec-36p::wrmScarlet*), *bus-1p::GFP*, pBJ36]; | This study | MBA1919 |
| **Oligonucleotides** |  |  |
| See **Table S2** for the list of oligonucleotides used |  |  |
| **Recombinant DNA** |  |  |
| Plasmid: *clec-35 3'UTR:wrmScarlet::clec-35/27p::GFPo::clec-27 3'UTR* (in pUC57) | Genewiz | pCV12 |
| Plasmid: *rab-3p::clec-35::T2A::clec-27::unc-54 3'UTR* (in pBJ97) | This study | pCV10 |
| Plasmid: *dpy-7p::clec-35::T2A::clec-27::unc-54 3'UTR* (in pUC57) | This study | pCV11 |
| Plasmid: *vha-6p::clec-35::T2A::clec-27::unc-54 3'UTR* (in pBJ97) | This study | pKL158 |
| Plasmid: *srg-8p::clec-27::clec-27 3'UTR* (in pBJ97) | This study | pKL1 |
| Plasmid: *gcy-15p::clec-27::clec-27 3'UTR* (in pBJ97) | This study | pKL2 |
| Plasmid: *gcy-33p::clec-27::clec-27 3'UTR* (in pBJ97) | This study | pKL3 |
| Plasmid: *str-1p::clec-27::clec-27 3'UTR* (in pBJ97) | This study | pKL4 |
| Plasmid: *ceh-37p::clec-27::clec-27 3'UTR* (in pBJ97) | This study | pKL6 |
| Plasmid: *odr-10p::clec-27::clec-27 3'UTR* (in pBJ97) | This study | pKL7 |
| Plasmid: *odr-7p::clec-35::T2A::clec-27::unc-54 3'UTR* (in pBJ97) | This study | pKL8 |
| Plasmid: *str-1::clec-35::T2A::clec-27::unc-54 3'UTR* (in pBJ97) | This study | pKL128 |
| Plasmid: *odr-10p::GFPo::unc-54 3'UTR* (in pBJ97) | This study | pKL53 |
| Plasmid: *str-1p::GFPo::unc-54 3'UTR* (in pBJ97) |  | pKL54 |
| Plasmid: *clec-36 3'UTR:wrmScarlet::clec-36/26p::GFPo::clec-26 3'UTR* (in pUC-GW-AMP) | Genewiz | N/A |
| Plasmid: *vha-6p::clec-36::T2A::clec-26::unc-54 3'UTR* (in pBJ97) | This study | pKL141 |
| Plasmid: *rab-3p::clec-36::T2A::clec-26::unc-54 3'UTR* (in pBJ97) | This study | pKL136 |
| Plasmid: *odr-7p::clec-36::T2A::clec-26::unc-54 3'UTR* (in pBJ97) | This study | pKL63 |
| Plasmid: *str-1p::clec-36::T2A::clec-26::unc-54 3'UTR* (in pBJ97) | This study | pKL130 |
| **Software and algorithms** |  |  |
| GraphPad Prism 7 | GraphPad Software, La Jolla, CA | https://www.graphpad.com/scientificsoftware/prism/ |
| Fiji | Schindelin et al., 2012<sup>75</sup> | https://imagej.net/software/fiji/ |
| Kallisto | Bray et al., 2016<sup>76</sup> | https://pachterlab.github.io/kallisto/ |
| OASIS 2 | Han et al., 2016<sup>77</sup> | https://sbi.postech.ac.kr/oasis2/ |
| MEGA 11 | Tamura et al., 2021<sup>78</sup> | https://www.megasoftware.net/ |

## RESOURCE AVAILABILITY

### Lead Contact

Further information and requests for resources and reagents should be directed to and will be fulfilled by the Lead Contact, Michalis Barkoulas.

### Materials Availability

All unique/stable reagents generated in this study are available from the Lead Contact without restriction.

### Data and Code Availability

The raw RNA-seq data have been deposited to NCBI GEO under accession GEO: GSE263858

## EXPERIMENTAL MODEL AND SUBJECT DETAILS

All *C. elegans* strains were cultured on NGM plates seeded with *E. coli* OP50, at 20°C under standard conditions as previously described^79^. Continuous cultures of oomycete isolates were maintained by chunking infected animals on an OP50-NGM plate whenever the plate was running out of food with intermittent addition of fresh N2^12,48^. The list of all the strains used in this study is available in the supplementary Table S2.

## METHOD DETAILS

### EMS mutagenesis

L4 stage animals carrying the *chil-27p::GFP* reporter were mutagenized with 24 mM ethyl methanesulfonate (EMS) (Sigma) in 4 mL M9, for 4 hours with intermittent mixing by inversion. Mutagenised animals were then washed 10 times with 15 mL M9 to completely get rid of EMS from the suspension and plated onto 90 mm NGM plates seeded with *E. coli* OP50. After brief recovery (4-12h), adults were randomly picked and divided into 90 mm plates carrying 10 animals in each. After 72 hours at 20°C, all F1 gravid adults in each plate were individually bleached and respective pool of F2 embryos were collected onto new plates containing *M. humicola* extract prepared as previously described^49^. Around 120,000 haploid genomes were screened in multiple independent screens to identify animals showing loss of *chil-27p::GFP* induction upon extract treatment. The causative mutations *ceh-37(icb36), clec-27(icb104),* and *clec-35(icb105)* were identified by crossing mutants to the polymorphic CB4856 strain, where 15-25 F2 recombinants selected for loss of *chil-27p::GFP* induction upon extract treatment were pooled in equal proportion to obtain genomic DNA for whole genome sequencing (Novogene, UK). WGS data was analysed using the CloudMap Hawaiian variant mapping pipeline to identify the causative mutations^80^. Additional alleles were identified by direct WGS of mutants without crossing to CB4856.

### Oomycete infection assays

Five *M. humicola* infected animals (filled with spherical sporangia) were added to the lawn of 100 µl *E. coli* OP50 on three 35 mm NGM plates and 20 live L4 animals for each genotype were transferred individually to each plate (n=60 per condition). Dead animals with visible sporangia were scored every 48 hours and live animals were transferred to a new NGM plate with five infected animals to maintain constant pathogen load throughout the experiment. Dead animals without evidence of infection or animals missing from the plate were censored. All infection assays were performed in triplicates at 20°C. Survival curves were plotted using OASIS 2 and statistical significance was evaluated using Log-Rank test.

### RNAseq

For transcriptome analysis, animals were first synchronized by bleaching and were treated with *M. humicola* extract for 4 hours at L4 stage along with untreated animals as control. After this they were collected for RNA extraction using TRIzol (Invitrogen) and isopropanol/ethanol precipitation. RNA was quantified using NanoDrop (Thermo Scientific) and quality was assessed by gel electrophoresis. The samples were collected from three independent biological repeats for each condition and RNA sequencing was completed by Novogene (UK). Kallisto was used for alignments with the WS283 transcriptome from Wormbase^76^. Count analysis was performed using Sleuth along with a Wald Test to calculate log_2_fold changes^81^. All RNAseq data files are publicly available from the NCBI GEO database under the accession number GSE263858.

### Microscopy

For performing induction assays with oomycete extracts, day 1 gravid adults were bleached, and the embryos obtained were plated onto standard NGM-OP50 plates at 20°C. Oomycete extracts from *M. humicola*^12^ and *H. zoospora*^48^ were made as previously described. Extract treatment was performed by adding 150-200 µl of extract on the NGM plate such that it fully covered the OP50 lawn. For most experiments, embryos were treated with extract for 48 hours and imaging of L4 stage animals was performed. For specific experiments as described in the results section, extract treatment was performed from L2 or L4 stage onwards and day 1 adults were imaged after 48 hours or 24 hours respectively. Animals to be imaged were picked into a drop (5µl) of M9 containing sodium azide (25 mM) on an agarose (2%) pad made on a glass slide. To document *chil-27p::GFP* expression, we used a Zeiss Axio Zoom V16 microscope and images were taken using the associated ZEN Microscopy software.

To image *clec-27p::GFP,* animals were mounted on a slide as described above, and were observed at 40x magnification on a Zeiss Compound microscope (AxioScope A1) with images taken using OCULAR (QImaging). To look at transcriptional reporters of *clec-27/35* and *clec-26/36* or co-localization of *clec-35p::mScarlet* and *clec-36p::mScarlet* with neuron-specific markers, confocal microscopy was performed on Leica SP8 inverted confocal microscope at 63x magnification, with excitation of GFP by an Argon 488nm laser and excitation of mScarlet by a He-Ne 594nm laser. To image CEH-37::GFP, L1 stage animals were observed at 100x magnification on Nikon Ti-eclipse epifluorescence microscope with an iKon M DU-934 CCD camera (Andor).

### Molecular cloning and transgenesis

For rescue of mutants obtained from the EMS screen, *clec-27* and *clec-35* were PCR amplified from N2 genomic DNA using primers P1 and P2, and P3 and P4 respectively. These were injected in the respective mutants at 20 ng/µl. For rescue of *ceh-37(icb36),* fosmid (WRM065bE05) carrying the WT copy of the gene was injected at 20 ng/µl. To generate clec-27/35 transcriptional reporter strains, a synthetic gene fragment (*clec-35 3’UTR:wrmscarlet::clec-35/27p::GFPo::clec-27 3’UTR*) cloned in pUC57 was obtained from Genewiz. The resulting plasmid, pCV12, was injected in N2 animals at 50 ng/µl. To generate a construct for pan-neuronal expression of *clec-27/35*, a fragment containing *clec-35* was PCR amplified from N2 genomic DNA using primers P5 and P6, a fragment containing *clec-27* was amplified from N2 genomic DNA using primers P7 and P8, a fragment containing T2A was amplified using primers P9 and P10, and all 3 fragments were then inserted into the vector pGO29 (*rab-3p::unc-54 3’UTR*) by Gibson cloning, generating pCV10(*rab-3p::clec-35::T2A::clec-27*). For epidermal rescue, the fragment *clec-35::T2A::clec-27* was PCR amplified from pCV12 using primers P11 and P12 and was inserted into vector pIR6 (*dpy-7p::unc-54 3’UTR*) by Gibson cloning, generating pCV11(*dpy-7p::clec-35::T2A::clec-27*). For intestinal rescue, the promoter for gene *vha-6* (3kb upstream of TSS) was PCR amplified from N2 genomic DNA using primers P13 and P14 and the vector backbone containing *clec-35::T2A::clec-27* was PCR amplified from pCV12 using primers P15 and P16. The two fragments were assembled by Gibson cloning to make pKL158(*vha-6p::clec-35::T2A::clec-27*). pCV10, pCV11 and pKL158 were injected in *clec-35(icb105)* mutant animals at 5 ng/µl. For neuronal and intestinal expression of *clec-27/35* in *ceh-37(icb36)* mutants, pCV10 and pKL158 were injected in *ceh-37(icb36)* mutant animals at 5 ng/µl. All microinjections included pRJM163 (*bus-1p::GFP*) as the co-injection marker at 25 ng/µl, along with pBJ36 as carrier DNA to bring the total DNA concentration of the mix to 110 ng/µl.

For neuron-specific rescue of *clec-27*, the fragment *clec-27::clec-27 3’UTR* was amplified using the primers P17 and P18, the promoters for neuron-specific expression were amplified using P19 and P20 for *ceh-37p* fragment, P21 and P22 for *odr-10p* fragment, P23 and P24 for *str-1p* fragment, P25 and P26 for *srg-8p* fragment, P27 and P28 for *gcy-33p* fragment, P29 and P30 for the *gcy-15p* fragment. Each neuron-specific promoter fragment was inserted along with the *clec-27::clec-27 3’UTR* fragment into SpeI digested pBJ97 by Gibson cloning, generating the plasmids pKL6(*ceh-37p::clec-27*), pKL7(*odr-10p::clec-27*), pKL4(*str-1p::clec-27*), pKL2(*gcy-15p::clec-27*), pKL3(*gcy-33p::clec-27*), and pKL1(*srg-8p::clec-27*). These plasmids were injected into *clec-27(icb104)* mutant animals at 50ng/µl, including pRJM163 (*bus-1p::GFP*) as the co-injection marker at 25 ng/µl, along with pBJ36 as carrier DNA to bring the total DNA concentration of the mix to 110 ng/µl.

For AWA specific expression of *clec-35/27*, the *clec-35::T2A::clec-27::unc-54* fragment was PCR amplified from pCV12 using primers P31 and P32. pKL7 was digested with AvrII, generating a backbone containing *odr-7p*, which was assembled by Gibson cloning with the *clec-35::T2A::clec-27::unc-54* fragment, generating pKL8(*odr-7p::clec-35::T2A::clec-27*). For AWB specific expression, the vector backbone containing *clec-35::T2A::clec-27::unc-54* was PCR amplified from pKL8 using primers P33 and P16. The *str-1p* fragment was amplified from pKL4 using primers P34 and P35. The two fragments were then assembled by Gibson cloning, generating pKL128(*str-1::clec-35::T2A::clec-27*). These plasmids were injected into *ceh-37(icb36)* mutant animals at 50ng/µl, including pRJM163 (*bus-1p::GFP*) as the co-injection marker at 25 ng/µl, along with pBJ36 as carrier DNA to bring the total DNA concentration of the mix to 110 ng/µl.

For the AWA GFP marker (*odr-10p::GFPo::unc-54*), the *odr-10p* fragment was amplified using primers P36 and P37. The GFPo fragment was amplified using primers P38 and P39. The *unc-54 3’ UTR* fragment was amplified using primers P40 and P41. Linearized vector was amplified using primers P42 and P16 from the pBJ97 plasmid. These three fragments were assembled with the vector by Gibson cloning, generating pKL53. For the AWB GFP marker (*str-1p::GFPo::unc-54*), the *str-1p* fragment was amplified using primers P34 and P43. The GFPo fragment was amplified using primers P38 and P39. The *unc-54 3’ UTR* fragment was amplified using primers P40 and P41. Linearized vector was amplified using primers P42 and P16 from the BJ97 plasmid. These three fragments were assembled with the vector by Gibson cloning, generating pKL54. These plasmids were injected into N2 animals at 40ng/µl, including PIP1303F (*rps-0::hgb::unc-54*) as the co-injection marker at 15 ng/µl, along with pBJ36 as carrier DNA to bring the total DNA concentration of the mix to 110 ng/µl.

The *clec-26/36* transcriptional reporter was designed with *wrmScarlet* replacing the *clec-36* RNA coding sequence and *gfp* replacing the *clec-26* RNA coding sequence, whilst keeping the *clec-26/36* shared promoter intact. This transcriptional reporter was commercially synthesized (Genewiz) in a pUC-GW-AMP vector. The plasmid was then injected into N2 animals at 50ng/µl, including pRJM163 (*bus-1p::GFP*) as the co-injection marker at 25 ng/µl, along with pBJ36 as carrier DNA to bring the total DNA concentration of the mix to 110 ng/µl.

For rescue of *clec-26/36* in different tissues, the *vha-6p* (intestinal) promoter fragment was amplified using primers P13 and P44, the *rab-3p* (neuronal) promoter fragment was amplified using primers P45 and P46, the *odr-7p* (AWA) promoter fragment was amplified using primers P47 and P48, and the str-1p (AWB) promoter fragment was amplified using P34 and P49. Each promoter fragment was combined with a linearized vector containing *unc-54 3’ UTR* (amplified using primers P50 and P16 from pKL53), a fragment containing *clec-36::T2A* (amplified from N2 gDNA using primers P51 and P52), and a fragment containing *T2A::clec-26* (amplified from N2 gDNA using primers P53 and P54), generating the plasmids pKL141(*vha-6p::clec-36::T2A::clec-26*), pKL136 (*rab-3p::clec-36::T2A::clec-26*), pKL63(*odr-7p::clec-36::T2A::clec-26*), and pKL130(*str-1p::clec-36::T2A::clec-26*). These plasmids were injected into *clec-26/36 (icb135)* mutant animals at 50ng/µl, including pRJM163 (*bus-1p::GFP*) as the co-injection marker at 25 ng/µl, along with pBJ36 as carrier DNA to bring the total DNA concentration of the mix to 110 ng/µl.

### CRISPR-mediated genome editing

To generate deletions in *clec-26*, *clec-36*, and both simultaneously, two crRNA were used for every deletion with one binding towards the 5’ end of the gene and the other towards the 3’ end. crRNA 1 and 2 used to make *clec-36(icb138)* deletion, crRNA 3 and 4 used to make *clec-26(icb130)* deletion, and crRNA 1 and 4 to delete both the genes *clec-26/36(icb135)* simultaneously. The sequences of all crRNA used can be found in Table S2 as well as the sequence/genomic coordinates of the deleted regions. For the injection mix, 1.4 µL of each crRNA (IDT) (from 34 µM stock) was mixed with 5 µL of tracrRNA (IDT) (from 18 µM stock) and 1 µL of Cas9 protein (IDT) (from 2.5 µg/µL stock) by pipetting several times before incubating at 37°C for 15 min. Following this, pRJM163 (*bus-1p::GFP*) was added at concentration of 25 ng/µl as co-injection marker and nuclease-free water was added to bring up the total volume to 20 µl. The mix was centrifuged at 13000g for 5 min and 17 µl was transferred to a new tube, which was the used for injecting the strain MBA281 (*C. elegans* carrying *chil-27p::GFP* reporter). Transgenic F1s were individually picked based on the presence of *bus-1p::GFP* spot in the tail and genotyping was performed using the following primer pairs: P55 crispr and P56 crispr for *clec-36(icb138),* P57 crispr and P58 for *clec-26(icb130),* P59 and P60 for *clec-26/36(icb135).* The sequences of all primers used can be found in Table S2.

### RNA Interference (RNAi)

RNAi by feeding was used as previously described^82^. The RNAi clone for *ceh-37* and *skn-1* were obtained from the ORFeome Library, while clones for all *clec* genes tested, *old-1, vab-3* and *pals-22* were obtained from the Ahringer Library (Source BioScience). For *flor-1* RNAi, a custom-made clone was used^55^. The clones were confirmed by sequencing prior to use. RNAi plates were made by adding 1 mM IPTG (Thermo scientific) to NGM to induce the expression of dsRNA along with 40 µg/ml ampicillin and 10 µg/ml tetracycline to select for the *E. coli* HT115 strain carrying the RNAi clones. For all RNAi treatments (except *skn-1* RNAi), 6 L4 stage animals (P0) were added onto RNAi plates and after 96 hours of incubation at 20°C the progeny at late L4 / early adult stage was scored. For *skn-1* RNAi, post-embryonic RNAi treatment was performed where embryos collected by bleaching gravid adults were added onto RNAi plates and day 1 adults after 72 hours of incubation at 20°C were scored for the phenotype. For experiments involving extract exposure and RNAi treatment, extract was added on the RNAi plates when the progeny was at L2 stage, 48 hours prior to scoring at the adult stage. All experiments were performed in biological triplicates, except the RNAi screen in Figure S4A, which was performed once with technical duplicates. At least 50 animals were scored for every condition in each repeat. Statistical significance was evaluated using chi-squared test.

### Single molecule fluorescence *in situ* hybridization

smFISH was performed as previously described^83^. Briefly, L2 stage N2 animals were fixed with 4% formaldehyde (Sigma-Aldrich) in 1x PBS (Ambion) for 45 min and were permeabilised with 70% ethanol for 24 hours. Hybridization was performed at 30°C for 16 hours. Probe sequences can be found in supplementary Table S2. Imaging was performed using the 100x objective in an inverted and fully motorised epifluorescence microscope (Nikon Ti-eclipse) with an iKon M DU-934 CCD camera (Andor) controlled via the NIS-Elements software (Nikon).

### Phylogenetic Analysis

To identify all the *clec* pairs in the *C. elegans* genome, 1 kb sequence upstream of TSS (predicted promoter region) for all *clec* genes was recovered using WormBase ParaSite followed by BLAST search against *C. elegans* genome. The sequences that aligned with any *clec* gene were identified and 36 *clec* genes present in pairs were identified within the *C. elegans* genome (*clec-128/129, clec-131/132, clec-62/63, clec-158/159, clec-167/168, clec-197/198, clec-73/74, clec-83/84, clec-45/46, clec-34/38, clec-29/39, clec-23/40, clec-26/36, clec-27/35, clec-22/31, clec-24/32, clec-245/246, clec-255/256*). Among these genes, 4 genes are annotated as pseudogenes (*clec-131, clec-159, clec-46, and clec-23*) on Wormbase and these pairs were eliminated from further analysis. Protein sequences of the remaining 28 paired protein-coding *clec* genes were aligned in MEGA11^78^ by MUSCLE and the resulting alignment was used to generate a phylogenetic tree by maximum-likelihood with 500 bootstrap replications.

### Heterologous *clec* expression and co-immunoprecipitation

Constructs for heterologous expression of various full length CLEC proteins were generated by Gibson assembly of a PCR fragment amplified from *C. elegans* cDNA into EcoRV linearised pKGC3S harbouring a C-terminal GFP-or myc-tag (fragments were amplified using the primer pairs P61/P62, P63/P64, P65/P66, P67,P68). *N. benthamiana* plants were grown in a controlled environment growth chamber with a temperature range of 22–25°C, humidity of 45–65% and a 16/8-h light/dark cycle. Three-week-old plants were grown in individual pots until infiltration with *Agrobacterium*. *A. tumefaciens* GV3101 was transformed with *clec* constructs and grown overnight in LB with appropriate antibiotics at 28°C. Bacteria were collected and resuspended in infiltration buffer (10 mM MES, 10 mM MgCl2, and 150 μM acetosyringone (pH 5.6) at a final OD_600_ of 0.5 for constructs and 0.05 for p19 in combinations as indicated. Leaf discs were collected 2 dpi and shock frozen until protein extraction. CLEC proteins were extracted by lysis with bead beating (2 min at 1500 rpm). 400 µl extraction buffer (10% (v/v) glycerol, 25 mM Tris, 1 mM EDTA, 150 mM NaCl at pH7.4 with 1 mM PMSF, 1 mM DTT, 0.3% NP-40) was added and tubes incubated on ice for 20 min and regular shaking. The leaf suspension was centrifuged for 10 min at 17,000 xg, the supernatant transferred and centrifuged again for 10 min at 17,000 xg. The supernatant was incubated for 1h at 4°C with equilibrated GFP and Myc beads (Proteintech) successively, and washed 3x with extraction buffer before proteins were eluted with SDS loading buffer at 70°C for 10 min. Samples were run on a NuPAGE™ 4 to 12%, Bis-Tris Mini gel and then transferred at 30V for 75 min to a nitrocellulose membrane using a Mini gel tank (Invitrogen). Membranes were blocked in 3% BSA in PBST (137 mM NaCl, 12 mM Phosphate, 2.7 mM KCl, pH 7.4, 0.1% tween-20) before adding the primary antibody at 1:1500 dilution (α-GFP, rabbit, Invitrogen; α-myc, rat, Proteintech) for incubation overnight at 4°C. The membrane was washed with PBST before adding the secondary HRP-coupled antibodies at 1:3000 dilution (α-rabbit, sigma) or at 1:3000 dilution (α-rat, Merck). Blots were visualised with ECL (Amersham) on a Licor Odyssey Fc Imaging System.

## QUANTIFICATION AND STATISTICAL ANALYSIS

Graphic representation and statistical analysis were performed using the GraphPad Prism 7 software. Data shown in bar graphs indicate mean, and error bars represent standard error of proportion, standard error of the mean, or standard deviation, as indicated in respective figure legends. A chi-squared test, unpaired t-test, or Cochran Mantel Haenszel test was used to evaluate significance as indicated in respective figure legends. Survival data was analysed using OASIS 2 (https://sbi.postech.ac.kr/oasis2/)^77^ and significance was assessed using a log-rank test. Results were considered statistically significant when p < 0.05. Asterisks in figures indicate corresponding statistical significance as it follows: * p < 0.05; ** p < 0.01; *** p < 0.001; **** p < 0.0001.

## ACKNOWLEDGEMENTS

This work was supported by the Wellcome Trust [219448/Z/19/Z] and BBSRC [BB/X001865/1]. We thank Mark Hintze for bioinformatic support and Jonathan Saunders for comments on the manuscript. Some strains were provided by the CGC, which is funded by NIH Office of Research Infrastructure Programs (P40 OD010440). We thank Javier Apfeld for providing the AWA ablation strain and Rachel McMullan for the pRJM163 plasmid (*bus-1p::GFP*).

## CONTRIBUTIONS

KL and MG conducted most experiments and analysed the data. FT performed the co-IP experiment. CV-P and DI performed the genetic screens. KL, MG, and MB wrote the manuscript. All authors edited and approved the final manuscript.

---

**Figure S1:**
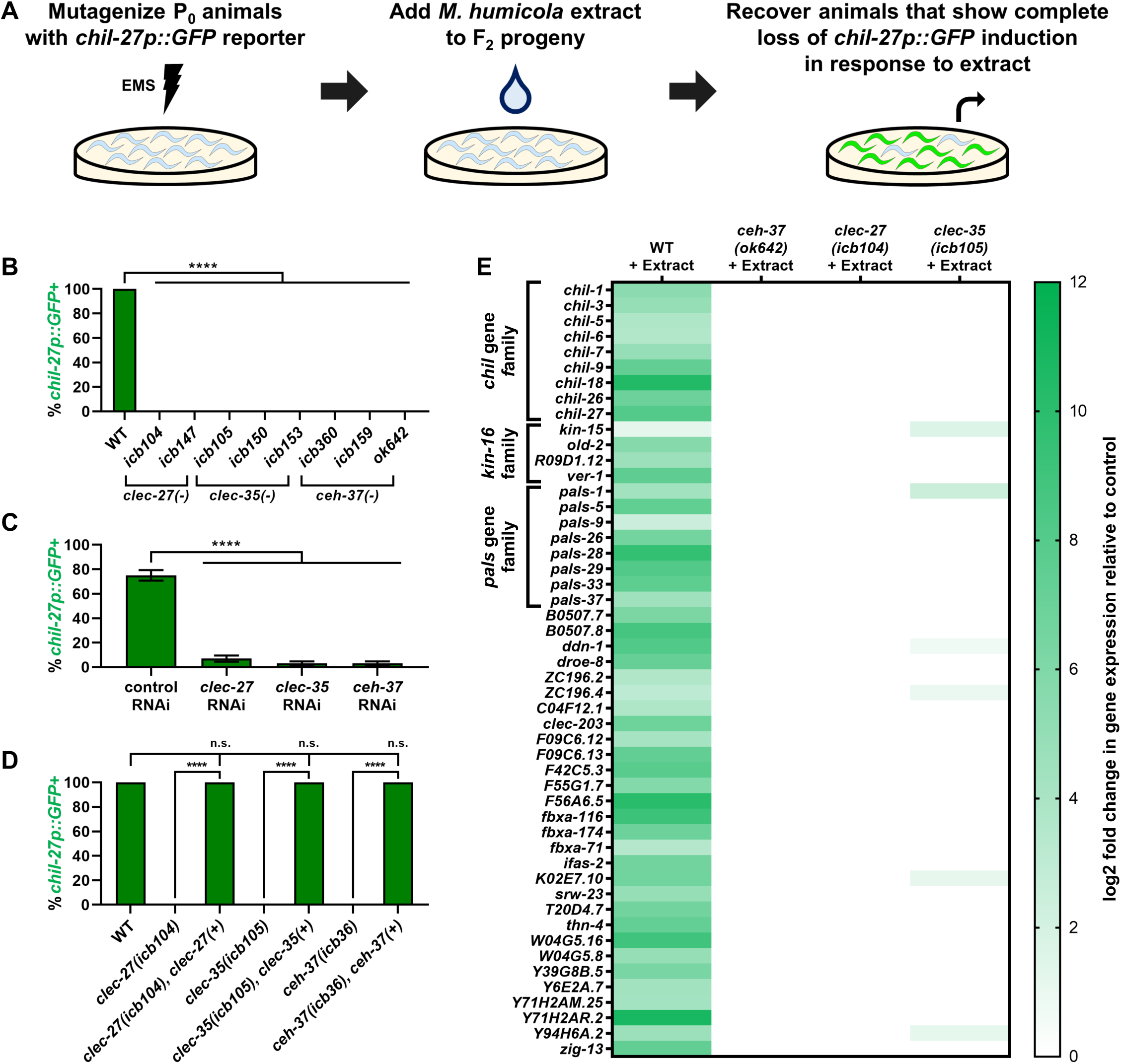
Validation of loss of ORR activation in mutants from the EMS screen. **(A)** Schematic of the forward genetic screen used in the study to recover mutants that do not respond to *M. humicola* extract exposure by inducing *chil-27p::GFP*. **(B)** *chil-27p::GFP* induction upon *M. humicola* extract exposure in all alleles shown in figure 1A (n=50 animals*, ****p<*0.0001 based on chi-squared test). **(C)** RNAi of *clec-27*, *clec-35* and *ceh-37* on wild-type animals strongly reduces *chil-27p::GFP* induction upon *M. humicola* extract treatment as compared to control RNAi (n=100, *\*\*\*\*p<*0.0001 based on chi-squared test). **(D)** Quantification of the genetic rescue of *clec-27(icb104), clec-35(icb105)* and *ceh-37(icb36)* shown in figure 1C (n=50 animals, *\*\*\*\*p<*0.0001 based on chi-squared test). **(E)** Heat map showing log2 fold change in expression of the 50 most reproducible ORR genes in wild-type animals, *clec-27(icb104), clec-35(icb105),* and *ceh-37(ok642)* mutants exposed to *M. humicola* extract in comparison to untreated control. log2 fold change is shown in green, while white represents nonsignificance. Note that all mutants show complete or near complete loss of ORR gene expression. Error bars in B, C, and D represent standard error of proportion.

**Figure S2:**
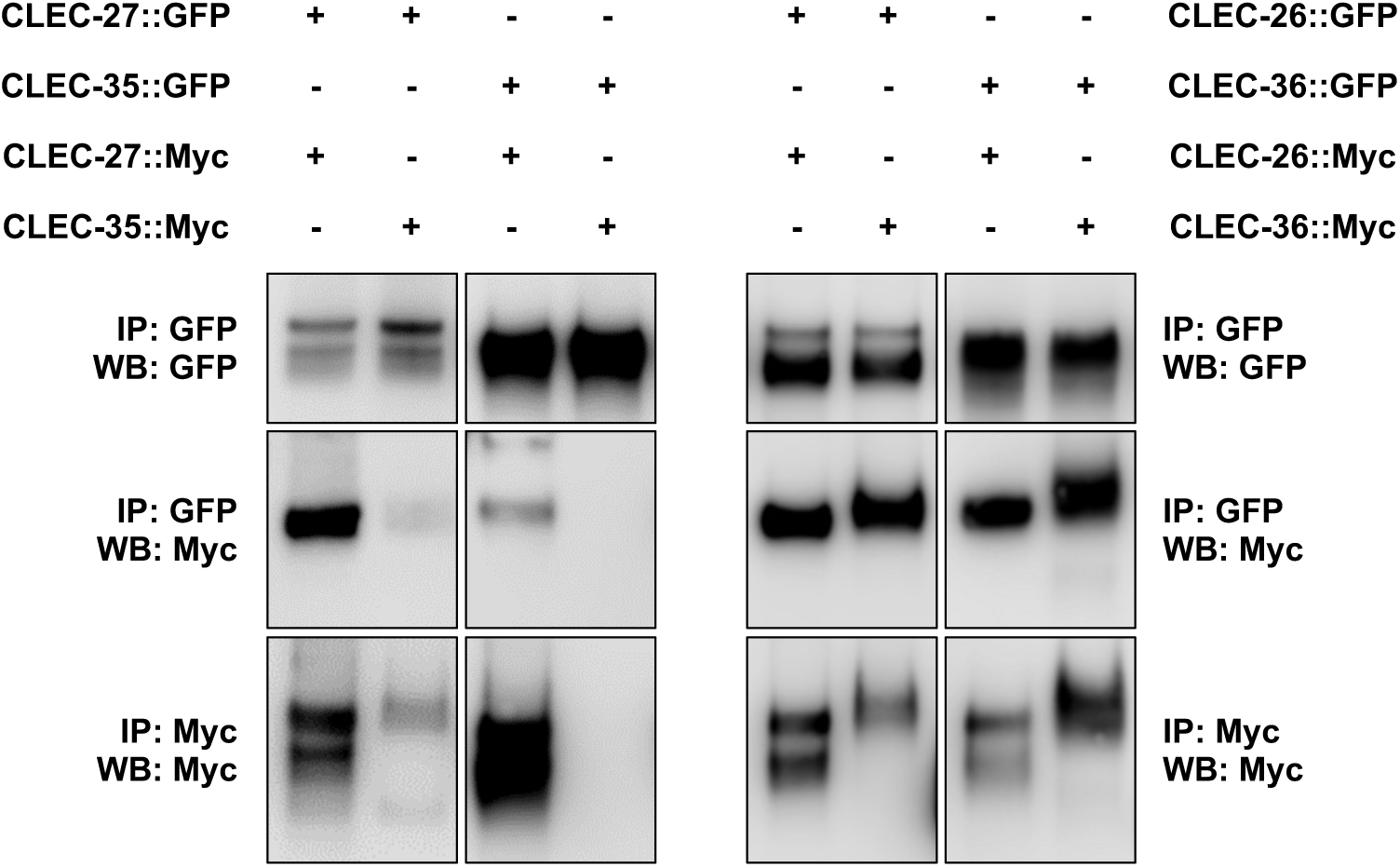
Interaction between CLEC proteins within pairs. Pull down experiments confirm the potential of interaction within the CLEC pairs (CLEC-27/35 on the left and CLEC-26/36 on the right). Note that due to the insolubility of CLEC proteins in prokaryotic expression systems and low expression levels in *C. elegans*, different combinations of GFP-and Myc-tagged CLEC proteins were expressed heterologously in *N. benthamiana* and extracted 2 days post infiltration for co-immunoprecipitation with GFP and Myc beads successively. The Myc IP serves as a loading control of Myc-tagged proteins because of low expression levels preventing reliable detection in input samples.

**Figure S3:**
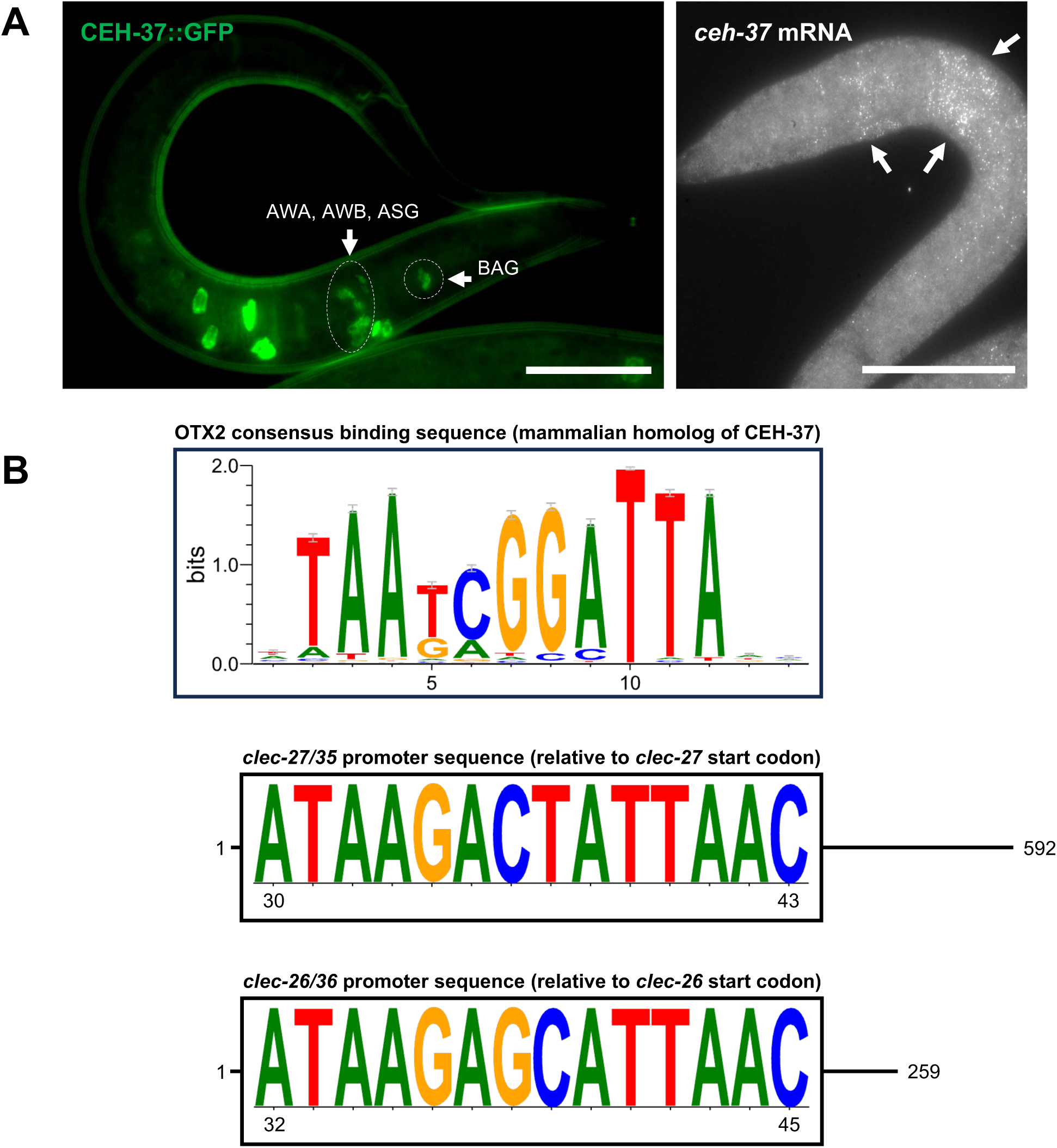
Expression of *ceh-37* and identification of putative binding sites in the *clec-27/35* and *clec-26/36* shared promoter. **(A)** CEH-37::GFP expression in specific foci around the pharynx where AWA, AWB, BAG and ASG neurons are located as well as in the anterior intestine (left panel). Right panel shows the expression of *ceh-37* mRNA by smFISH in anterior intestine as well as the head region. Scale bar for left panel is 20 µm and for right panel is 10 µm. **(B)** A putative binding motif for CEH-37 based on the mammalian homolog OTX2 is present in approximately the same region within the *clec-27/35* and *clec-26/36* shared promoters (*p*-value 5.94e-04 based on Tomtom comparison against a vertebrate database of known motifs).

**Figure S4:**
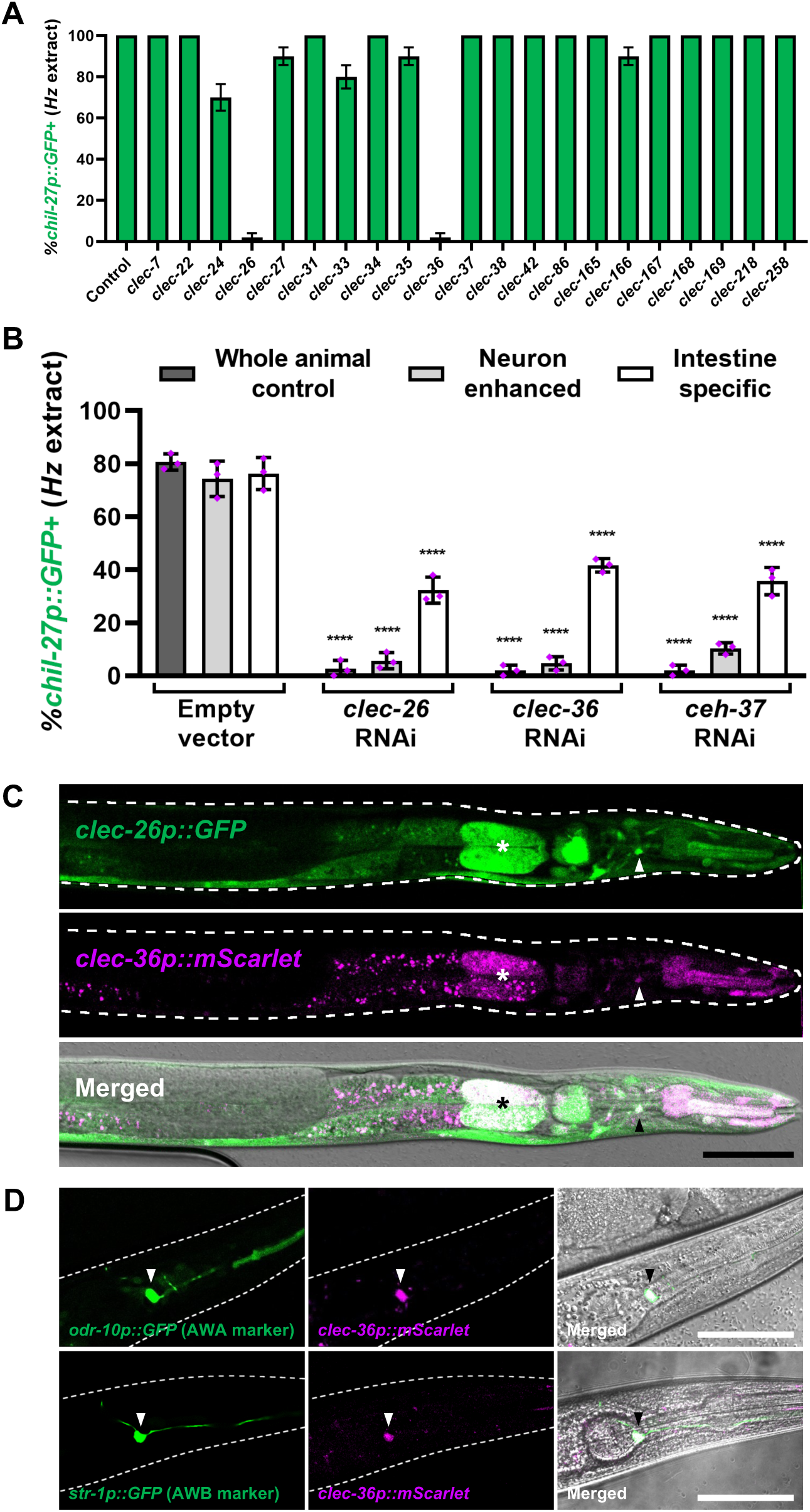
A CLEC-26/36 pair acts in the intestine and sensory neurons to mediate recognition of the oomycete *Haptoglossa zoospora*. **(A)** *chil-27p::GFP* induction upon *H. zoospora* extract treatment in wild-type animals treated with RNAi targeting CEH-37-regulated *clec* genes (n=50). **(B)** *chil-27p::GFP* induction upon *H. zoospora* extract exposure following tissue-specific RNAi of *clec-26, clec-36* and *ceh-37* (n=50 per condition, performed in triplicate, *\*\*\*\*p<*0.0001 based on chi-squared test). **(C)** Transcriptional reporter (*mScarlet::clec-26/36p::GFP*) made using the shared promoter of *clec-26/36* genes shows co-localization of GFP and mScarlet in neurons (marked with arrows) and anterior intestine (marked with asterisk). **(D)** Co-localization of *clec-36p::mScarlet* with AWA (*odr-10p::GFP*) and AWB (*str-1p::GFP*) specific markers confirms expression of *clec-26/36* in these neurons (shown with arrows). Scale bars in C and D are 50 µm. Error bars in A and B represent standard error of proportion.

